# Dopamine replacement remediates risk aversion in Parkinson’s disease in a value-independent manner

**DOI:** 10.1101/625467

**Authors:** Mariya V. Cherkasova, Jeffrey C. Corrow, Alisdair Taylor, Shanna C. Yeung, Jacob L. Stubbs, Martin J. McKeown, Silke Appel-Cresswell, A. Jon Stoessl, Jason J. S. Barton

## Abstract

**Introduction:** Clinical evidence suggests that Parkinson’s Disease (PD) patients are risk averse. Dopaminergic therapy has been reported to increase risk tolerance, but the underlying mechanisms are unclear. Some studies have suggested an amplification of subjective reward value, consistent with the role of dopamine in reward value coding. Others have reported value-independent risk enhancement. We evaluated the value-dependence of the effects of PD and its therapy on risk using tasks designed to sensitively measure risk over a wide range of expected values.

**Method:** 36 patients with idiopathic PD receiving levodopa monotherapy and 36 healthy matched controls performed two behavioural economic tasks aimed at quantifying 1) risk tolerance/ aversion in the gain frame and 2) valuation of potential gains relative to losses. PD patients performed the tasks on and off their usual dose of levodopa in randomized order; controls performed the same tasks twice.

**Results:** Relative to the controls, unmedicated PD patients showed significant value-independent risk aversion in the gain frame, which was normalized by levodopa. PD patients did not differ from controls in their valuation of gains relative to losses. However, across both tasks and regardless of medication, choices of the patients were more determined by expected values of the prospects than those of controls.

**Conclusion:** Dopamine deficiency in PD was associated with risk aversion, and levodopa promoted riskier choice in a value-independent manner. PD patients also showed an increased sensitivity to expected value, which was independent of levodopa and does not appear to result directly from dopamine deficiency.

## INTRODUCTION

Clinical evidence suggests that drug-naïve Parkinson’s disease (PD) patients are rigid and risk-averse [1,2]. On the other hand, dopaminergic therapy, particularly direct dopamine agonists, can cause impulse control disorders with compulsive risky reward seeking behaviors such as pathological gambling [3].

Pathological gambling is a form of irrational risk that disregards the expected value of outcomes; it is common knowledge that “the house always wins”. Given dopamine’s role in value coding, PD and its therapy could disrupt the guiding influences of expected value on choice, thereby encouraging irrational choices. Although a number studies using neurocognitive or behavioural economic decision-making tasks have suggested that dopaminergic therapy in PD is associated is with an increased willingness to take risks [4][5][6] [7][8][9][10], the findings have been mixed [11][12][13][14] and inconclusive with respect to the dependence of these effects on expected value. The latter can be gauged by modeling risky choice as a function of gamble characteristics in behavioural economic tasks. In two such studies, medication promoted riskier choice by amplifying the subjective value of rewards [7][8]. Another suggested it did so in a value-independent manner [9].

The types of gambles employed by the tasks may be relevant to these discrepancies: some studies used pure gamble tasks, where alternatives had the same valence [7][14][10], whereas others used mixed gambles pitting a certain outcome against a risky option that could result in either a gain or a loss [8][9]. The role of dopamine in coding the value of aversive events or punishments is less clear than in coding values of rewards [15]. Therefore, the requirement to evaluate potential losses alongside gains could obscure value-dependent effects in mixed gamble studies. The effects of PD and its therapy on valuation of gains relative to losses may be subtle: studies examining loss aversion – a tendency to avoid losses more than to seek equivalent size gains – have not produced clear cut findings, instead suggesting complex effects, possibly modulated by a history of depression [9][10].

We sought to evaluate the value-dependence of the risk-altering effects of PD and its therapy. To this end, we tested PD patients treated with levodopa monotherapy in ON and OFF states, as well as matched controls, using two behavioural economic tasks designed to probe risk attitudes as a function of the expected value. A two-choice lottery Vancouver Gambling Task [16][14] was used to assess willingness to take risks in the gain frame without the possibility of loss. To measure valuation of potential gains relative to losses, we used the Vancouver Roulette Task featuring mixed gambles with varying bet sizes.

We hypothesized that relative to controls, patients in the OFF state would demonstrate risk aversion both in the gain frame and in the context of mixed gambles, and that these risk-averse tendencies would be normalized by levodopa in a value-dependent manner.

## MATERIALS AND METHODS

### Participants

We tested 36 mildly to moderately affected patients with idiopathic PD and 36 age-matched (±5 years) healthy controls (Table 1). Patients were recruited from the Movement Disorders clinic at the University of British Columbia. Controls were spouses, friends and family members of the patients. The study was approved by the institutional research ethics board and carried out in accordance with the Declaration of Helsinki. Participants gave written informed consent.

**Table 1:** Participant demographics and clinical characteristics

|  | **PD (n=35)** | **Controls (n=35)** | **p** |
| --- | --- | --- | --- |
| **Age** | 64.83 (10.73) | 65.11(9.59) | 0.91 |
| **Gender (m/f)** | 26/9 | 21/14 | 0.20 |
| **Years of education** | 14.31 (2.92) | 14.42 (2.89) | 0.87 |
| **Right handed** | 31/35 | 30/35 | 0.72 |
| **PD duration** | 5.23 (3.96) | - | - |
| **Hoehn & Yahr** | 1.5 (0.61) | - | - |
| **UPDRS III** | 18.4 (10.8) | - | - |
| **LEED (mg)** | 646.42 (273.74) | - | - |
| **BDI** | 6.2 (3.45) | 4.24 (4.16) | 0.03* |
| **MoCA** | 27.4 (1.77) | 27.22 (1.73) | 0.68 |
| **CPGI** | 0.29 (.79) | 0.94 (2.02) | 0.08 |
| **TCI** | 7.08 (3.58) | 7.86 (2.46) | 0.29 |
| **QUIP-Short (n)** |  |  |  |
| Compulsive |  |  | 1 |
| Gambling | 0 | 0 |  |
| Compulsive sex | 1 | 0 | 0.31 |
| Compulsive eating | 1 | 1 | 1 |
| Compulsive buying | 0 | 1 | 0.31 |
| Hobbyism | 2 | 3 | 0.64 |
| Punding | 1 | 1 | 1 |

Unless otherwise specified, data are presented as mean (standard deviation). BDI: Beck Depression Inventory; MOCA: Montreal Cognitive Assessment; UPDRS III: Unified Parkinson's Disease Rating Scale, motor; CPGI: Canadian Problem Gambling Index; LEED: levodopa equivalent dose; QUIP: Questionnaire for Impulsive-Compulsive Disorders in Parkinson's Disease; TCI: Temperament and Character Inventory (TCI) [30]; Questionnaire for Impulsive-Compulsive Disorders in Parkinson's Disease-Rating Scale (QUIP) [29]. The Unified PD Rating Scale (UPDRS-III) scores were taken from the clinic database; the mean interval between the date of testing and UPDRS-III administration was $403 \pm 512.4$ days.

Exclusion criteria for both patients and controls were: other central neurological disorders, Montreal Cognitive Assessment [17] score < 24 OFF medication, Beck Depression Inventory [18] scores > 14, or ongoing treatment with antidepressants. One PD patient treated with a low dose of ropinirole (1mg daily) and one control participant with problematic gambling as per the Canadian Problem Gambling Index [19] were excluded from the analyses.

### Procedure

Patients were randomly assigned to one of two testing orders: 1) first session OFF, second session ON or 2) first ON, second OFF. For the OFF session, medication was withheld overnight, with the last dose at least 12 hours prior to the experiment for immediate release and at least 18 hours for controlled release levodopa. For the analyses, each control was matched to one PD patient, and their two testing sessions were given ON and OFF labels matching those of the PD patient, even though controls did not receive levodopa.

For the tasks described below, participants played to earn cash bonuses. The tasks were programmed in Experiment Builder (SR Research Ltd, Kanata, ON), and their order was pseudorandomized and matched between patients and controls.

#### Vancouver Gambling Task

On each trial, participants chose between two prospects: one featuring a larger and less probable gain and the other featuring a smaller and more probable gain. There were 10 different prospect pairs, each repeated in 10 trials. The difference in expected value (i.e. probability x magnitude of reward) between the two prospects in each pair ranged widely, from pairs that highly favored the smaller more probable prospect (the safer option) to pairs that highly favored the larger less probable prospect (the riskier option), as well as pairs that were close in expected value. The probability of gain for the two prospects always added up to one (0.2 / 0.8; 0.3 / 0.7; and 0.4 / 0.6), and reward magnitudes ranged from one to five tokens 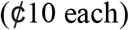. Thus, each pair had an Expected Value Ratio (EVR) computed as EV_(safe)_ – EV_(risky)_/mean(EV_(safe)_, EV_(risky)_) [14]. After each choice, the program determined the outcome of the wager based on its stated probability, and participants were given feedback as shown in Figure 1A.

**Figure 1:**
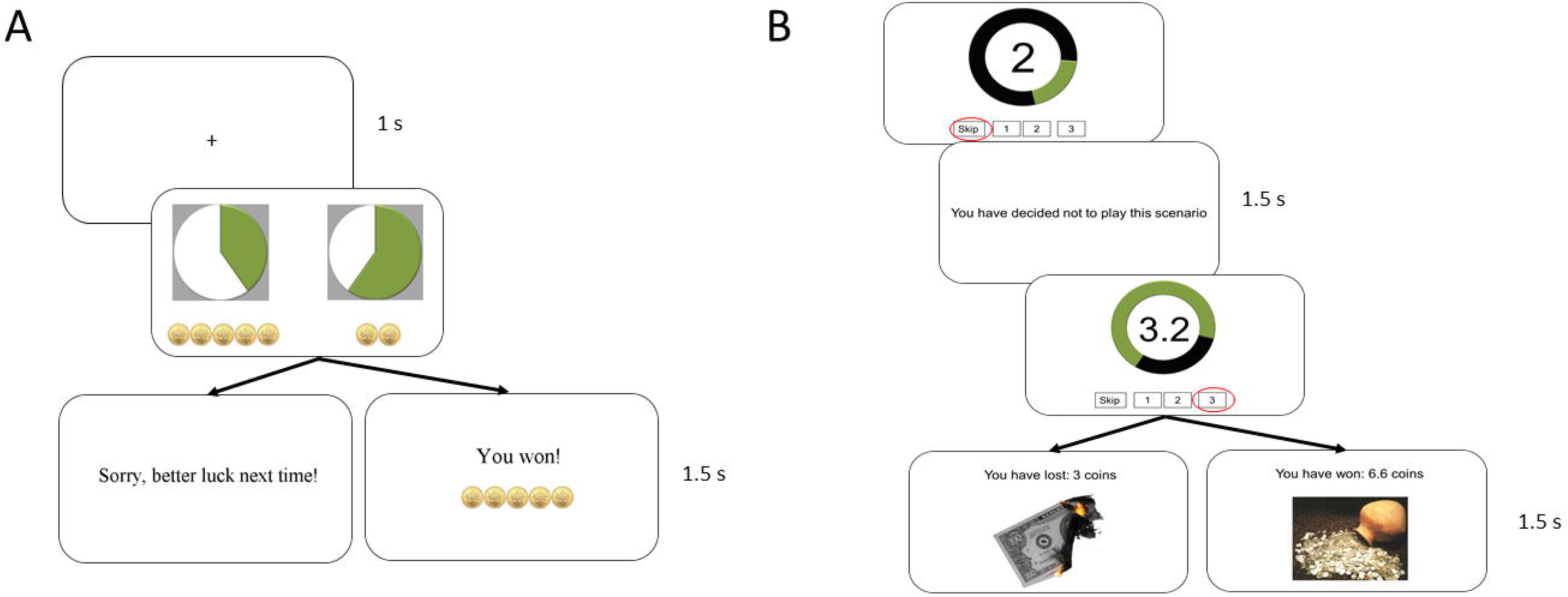
A) Vancouver Gambling Task (VGT); B) Vancouver Roulette Task (VRT). On the VRT, the green portion of the ring represents the probability of winning, the black portion represents the probability of losing, and the number in the centre of the ring is the multiplier.

#### Vancouver Roulette Task

This assessed the valuation of gains versus losses by asking participants to either accept or reject mixed gambles (Figure 1B). The gambles differed in the balance of the probability of gain versus the probability of loss, which always added up to one. If the participant accepted the gamble, they could bet one, two or three tokens. In the case of a win, the participant received the marginal return on the bet multiplied by a factor that also varied across trials (marginal return = bet*multiplying factor – bet). In the case of a loss, the participant lost the amount they bet. There were 17 different prospects, whose expected values were determined by the probability of gain versus loss and the size of the multiplier and ranged from highly favoring the gamble’s acceptance to highly favoring its rejection. Participants completed five blocks of 23 trials, for a total of 115 trials. Participants were given $10 at the start to offset possible losses.

Participants additionally performed a task of decision making under ambiguity as part of a larger study, which we report in the Supplement.

### Statistical Analyses

Demographic and clinical characteristics were analyzed using independent t-tests for continuous and chi square tests for categorical variables. The main analyses were performed using linear mixed effects models implemented via the lme4 package in R [20].

For the Vancouver Gambling Task, we first used a linear mixed effects model (glmer function) with a logistic link to model safe vs. risky choice on a trial-by-trial basis as a function of medication (ON vs. OFF or corresponding sessions in controls) in interaction with group. Random intercepts were modeled for participants, and random slopes were modeled for the effect of medication.

We then examined the effect of medication in the PD group alone and in interaction with gamble characteristics (EVR, probabilities and magnitudes). Because probabilities and magnitudes of the two alternatives were evaluated relative to each other, we performed isometric log ratio transformations to derive a single value for each. These relative probability and magnitude values were modeled as fixed effect terms in interaction with medication. The random effect structure was the same as above.

Finally, we ran separate models for the OFF and ON states to compare the likelihood of risky choice as a function of gamble characteristics between PD and control participants. EVR, probability and magnitude were modelled as fixed effects, and random intercepts were included for participants.

For the Vancouver Roulette Task, we modelled three variables describing participants’ choices: 1) accepting vs. rejecting a gamble; 2) betting 2 or more tokens; 3) betting 3 tokens. These decisions were modelled on a trial-by-trial basis using linear mixed effects models with a logistic link, first as a function of group and medication, then using separate models in PD and controls to examine the effects of gamble characteristics in interaction with medication, as well as models examining the effects of these characteristics in interaction with group. The random effect structure was same as with the other task.

Gender, age and task order were initially included as terms in all the models but were subsequently dropped as they did not significantly predict any outcomes of interest or improve the fit of the models.

## RESULTS

### Participant Characteristics

Patients and controls did not differ on demographic characteristics (Table 1). Although none of the participants were clinically depressed, PD patients had significantly higher BDI scores (p=0.03).

### Vancouver Gambling Task

There was a significant interaction between medication and group that predicted the likelihood of choosing the safer versus the riskier prospect (b = 0.58, SE = 0.26, z= 2.27, p = 0.02). The interaction was due to a difference in risk-taking between controls and PD patients in the OFF state but not the ON state (Figure 2). While the choices of PD patients in the ON state did not significantly differ from those of controls (p = 0.78), patients in the OFF state were more risk-averse than controls (main effect of group: b = 0.66, SE = 0.28, z= 2.32, p = 0.02). As expected, controls showed no change in performance from the OFF-corresponding session to the ON-corresponding session (p = 0.41). However, PD patients were significantly more likely to take risks in the ON than the OFF state (main effect of medication: b = 0.45, SE = 0.22, z= 2.08, p = 0.04). This medication effect did not interact with EVR, probability or magnitude (p_s_ ≥ 0.11).

**Figure 2:**
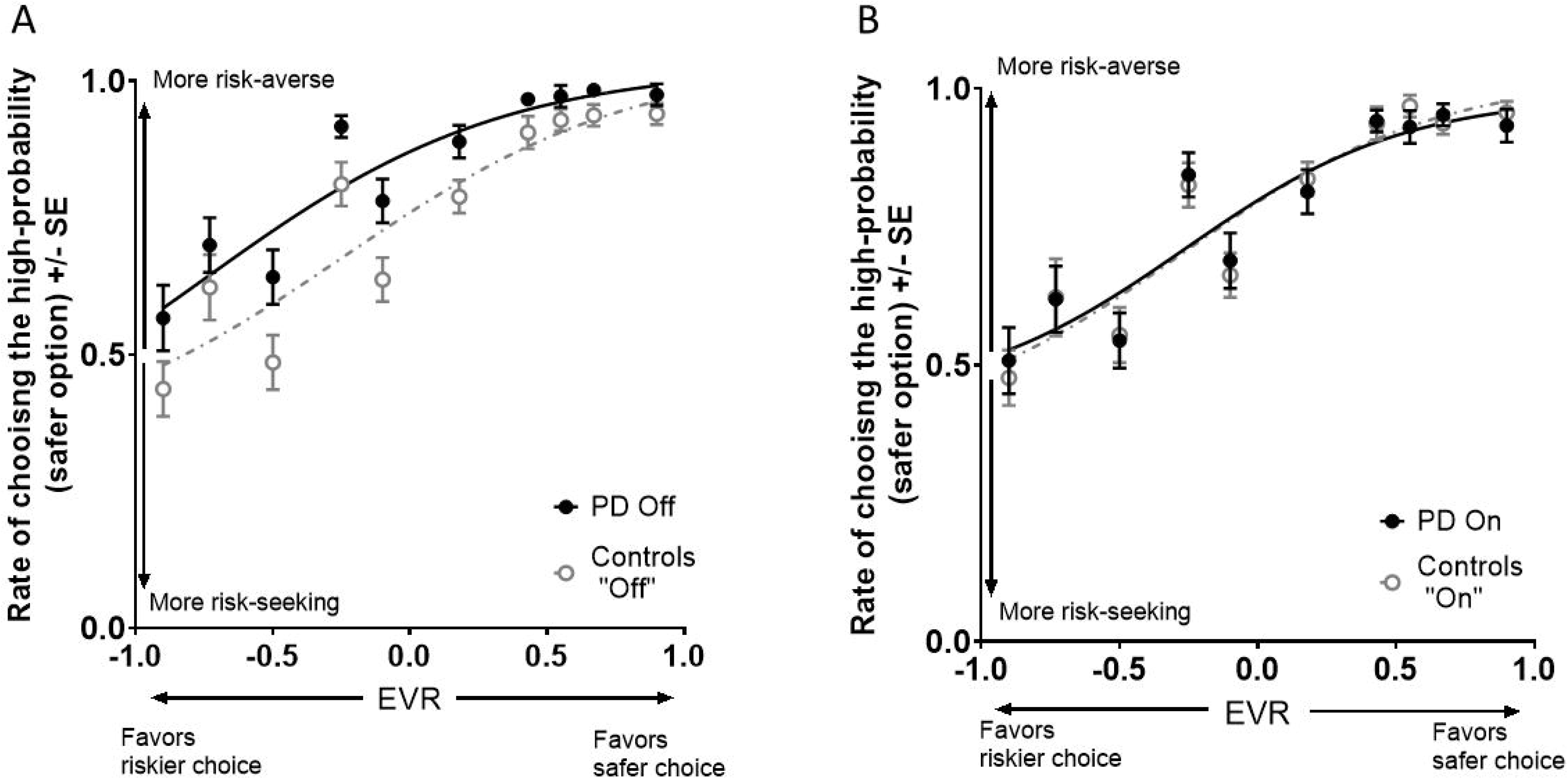
A) Vancouver Gambling Task performance off medication: PD vs. controls; B) VGT performance on medication PD vs. controls. Curves are fitted using a 4-parameter logistic function.

There were, however, significant interactions of group with gamble characteristics collapsing across the medication states. The interactions indicated that the patients’ choices were more driven than those of controls by the gambles’ probabilities (b = 1.03, SE = 0.27, z= 3.81, p = 0.0001); magnitudes (b = 0.52, SE = 0.15, z= 3.44, p = 0.0006); and expected values (b = 0.41, SE = 0.01, z= 4.09, p <0.0005).

### Vancouver Roulette Task

The interaction of group and medication state did not significantly predict the likelihood of accepting the gamble (p = 0.11), nor was there a main effect of group (p=0.29). Both controls and PD patients, whether in the ON or OFF state, disadvantageously accepted gambles with expected values < 0, in which losses were more likely or larger than gains (Figure 3 A&B). PD patients and controls also did not differ in their willingness to place higher bets of 2 or 3 tokens in either the OFF or ON state (p_s_ ≥ 0.21).

**Figure 3:**
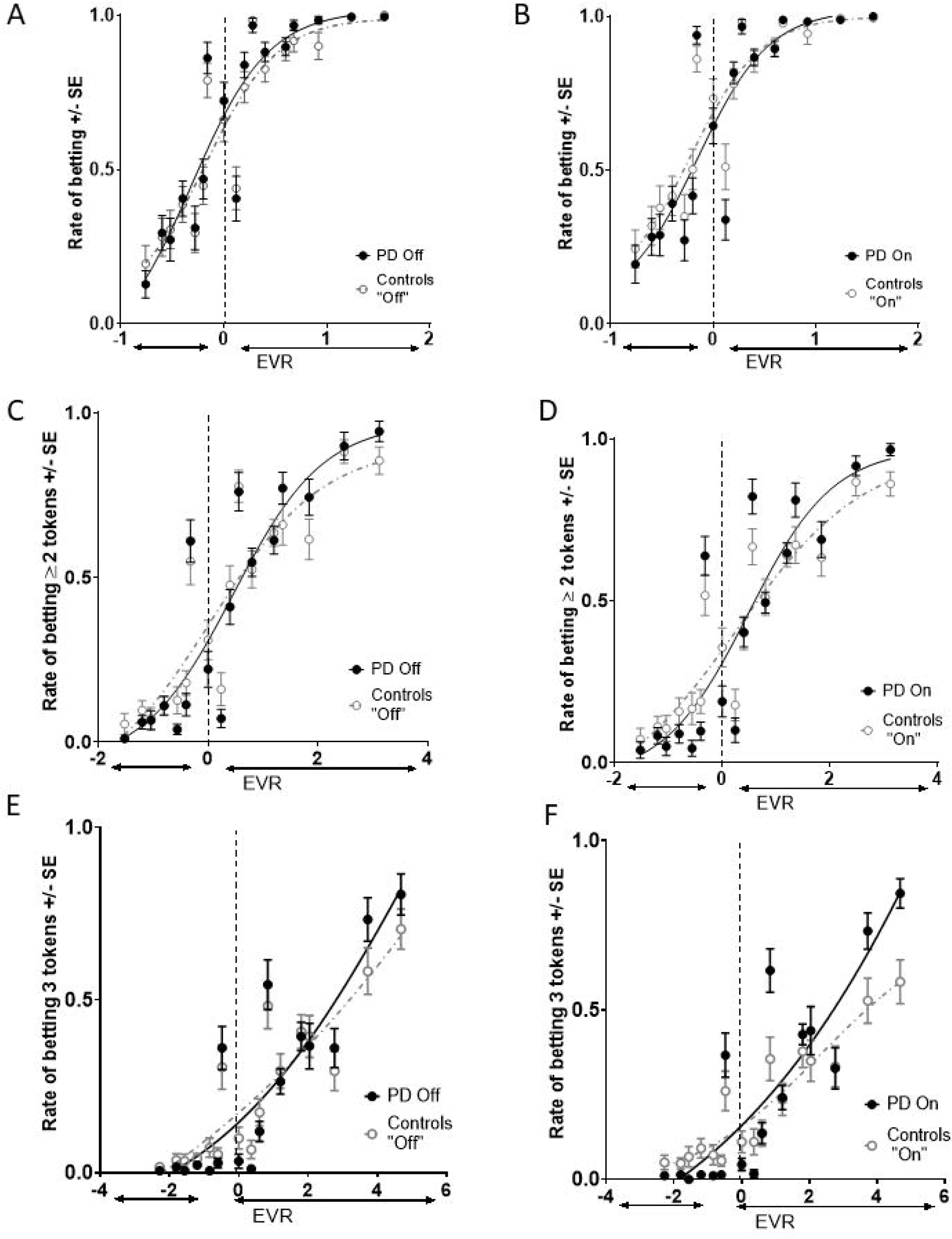
Rate of betting at least 1 token on the Vancouver Roulette Task as a function of the expected value (EV) of betting 1 token for PDs in the OFF state vs. controls (A) and PDs in ON state vs. controls (B). Rate of betting at least 2 tokens on the VRT as a function of the expected value (EV) for PDs in the OFF state vs. controls (C) and PDs in ON state vs. controls (D). Rate of betting 3 tokens on the VRT as a function of the expected value (EV) for PDs in the OFF state vs. controls (E) and PDs in ON state vs. controls (F).

However, again, relative to the controls, the patients’ likelihood of betting was more driven by prospect characteristics, as evidenced by a significant interaction of group with gain/ loss probability (b = 2.30, SE = 0.55, z=4.23, p < 0.0005), size of the multiplier (b = 1.1, SE = 0.04, z=25.35, p < 0.0005) and the gamble’s expected value computed as the difference between the EV of gain and the EV of loss (b = 4.2, SE = 0.11, z=39.81, p < 0.0005). This was also the case for the likelihood of placing higher bets: there was a significant interaction of group with gain/loss probability (≥2 tokens: b = 4.16, SE = 0.39, z=10.56, p < 0.0005; 3 tokens: b = 6.16, SE = 0.45, z=13.53, p < 0.0005) and the gamble’s expected value (≥2 tokens: b = 0.24, SE = 0.05, z=4.91, p < 0.0005; 3 tokens: b = 0.21, SE = 0.03, z=6.26, p < 0.0005), though not the size of the multiplier. This is reflected in the steeper slope of the patients’ betting likelihood function (Figure 3).

In addition, there was a significant main effect of “medication” collapsing between the groups (b = 0.29, SE = 0.14, z=2.05, p < 0.04), which was driven by the effect of testing session in controls (b = 0.31, SE = 0.14, z=2.25, p < 0.02); this effect was not significant in the patient group. An exploratory analysis showed that this effect in controls was not due to a practice effect (e.g. participants being more risk tolerant or averse in session 1 versus session 2; p=0.99).

In summary, PD patients were risk-averse OFF medication in the gain frame, but not in the context of mixed gambles. Levodopa normalized their risk aversion in a value-independent manner in the gain frame, resulting in more risk tolerant decisions. Regardless of the medication, patients’ choices were more strongly determined by the expected values of the prospects across both tasks.

## DISCUSSION

We found that unmedicated PD patients were risk averse for gain only gambles relative to matched controls. Although risk aversion in PD is often assumed, clear experimental evidence of it is surprisingly sparse. To our knowledge, only one prior study had clearly shown that PD patients were risk-averse in the OFF state relative to controls [8], although only a partial OFF state was achieved, in which levodopa but not dopamine agonists or other antiparkinsonian medications were withdrawn.

The current findings are at odds with those using an earlier version of the Vancouver Gambling Task [14], which failed to find OFF state risk aversion in the gain frame, though it did find risk aversion in the loss frame: patients showed a stronger preference than controls for less likely larger losses over smaller more likely ones. Differences in the representation of prospects may explain the discrepancy. Unlike the current study, which represented probabilities using pie charts, Sharp et al. represented probabilities numerically. Because neurotypical adults have difficulties with numerical ratio concepts such as probabilities [21], numeracy challenges may have masked risk aversion of PD patients in the gain frame, which was perhaps more apparent in the loss frame, as unmedicated patients are more sensitive to punishments [22]. Representation of prospects in decision-making tasks has been largely ignored and may deserve greater attention.

The patients’ risk aversion was normalized by levodopa in a value-independent manner, which echoes a finding in mixed sample of patients treated with levodopa and/ or dopaminergic agonists performing a mixed gamble task [9]. These results are in keeping with levodopa increasing value-independent gambling propensity in healthy volunteers [23][24], which was speculated to reflect a global potentiation of exploration-driven approach behaviour that could yield information gain from surprising outcomes. Other studies, however, suggested value-dependent effects [7][8], though one of the studies was geared towards detecting value-dependent, but not value-independent effects [8]. The absence of value-dependent effects in our study cannot be attributed to the potentially obscuring effects of loss prospects in mixed gambles, and it is unlikely that our task design prevented detection of value-dependent effects, as our tasks were optimized for their evaluation. It is possible, however, that reward magnitudes in our task were not sufficiently large to permit the expression of value-dependent effects, as utility function tends to roughly linear at smaller magnitudes.

We observed risk aversion in the PD patients and normalization thereof by levodopa only in the context of gains; valuation of potential losses relative to gains was unaffected by PD or levodopa. This is in keeping with previous studies suggesting unaltered loss aversion in PD [10][14][17], however the absence or risk aversion in the context of mixed gambles is somewhat surprising. This may be an artifact of the task and a possible limitation of the study: both patients and controls were biased in favor of betting even when the expected value of doing so was unfavorable. This bias may have resulted from either a) a house money effect, as participants were given $10 to play with at the outset, b) possible concern that frequent skipping of trials may be construed as poor participation or c) a framing effect, in which skipped trials may have been viewed as missed opportunities.

Despite our value-independent effects on risk, we observed that the decisions of the patients across both tasks were more strongly driven by expected value, while the choices of controls were more stochastic. This, to our knowledge, is a novel finding. The neurobiological substrates are unclear, as this was observed independent of levodopa. A plausible mechanism could be PD-driven adaptations at the level of postsynaptic dopamine signaling or other neurotransmitter systems. For example, there is some evidence of an upregulation of D2 receptors in early PD [26], which could influence the competition between the direct and the indirect basal ganglia pathways in determining choice, as could altered receptor sensitivity [27]. Indeed, a D2-specific pharmacological challenge in healthy volunteers resulted in less stochastic and more value-dependent choice [28]. At the cognitive level, impairments in reinforcement learning, which are well-documented in PD [22], could have contributed to the increased value-dependence of choice. A supplemental exploratory analysis of the effects of feedback history (streaks of wins and losses) on choice suggested that choices of controls tended to be more biased by feedback history effects, perhaps owing to implicit reinforcement learning, although this could not fully account for their choices being more stochastic (see Supplement).

One important limitation of our study is the absence of ON and OFF state UPDRS-III ratings. The ratings provided here were taken from our clinic’s database and were dated for some of the participants, so the disease staging data given in Table 1 may not be entirely accurate. In addition, we observed a significant difference between the “ON” and “OFF” session performance of the mixed gamble task in controls, which could indicate the task’s imperfect robustness.

In conclusion, we found that 1) PD patients were risk-averse for gains, but not when evaluating gains against losses; 2) their risk aversion likely resulted from dopamine deficiency, as it is normalized by levodopa; 3) regardless of medication, patients’ decision making was more value-driven, which might not be a direct result of dopamine deficiency, but could be a consequence of other PD-related neurobiological adaptations.

## Authors’ Roles

Mariya V. Cherkasova: project execution, statistical analysis, manuscript writing

Jeffrey C. Corrow: project execution, manuscript writing

Alisdair Taylor: project organization and execution

Shanna C. Yeung: project execution

Jacob L. Stubbs: project execution and manuscript review and critique

A. Jon Stoessl: project organization and manuscript review and critique

Martin J. McKeown: project organization and manuscript review and critique

Silke Appel-Creswell: project organization

Jason J. S. Barton: project conception, organization, statistical analysis review, manuscript review and critique

## REFERENCES

[1] M. Poletti, U. Bonuccelli, Personality traits in patients with Parkinson’s disease: Assessment and clinical implications, J. Neurol. 259 (2012) 1029–1038. doi:10.1007/s00415-011-6302-8.

[2] A. Dagher, T.W. Robbins, Personality, Addiction, Dopamine: Insights from Parkinson’s Disease, Neuron. 61 (2009) 502–510. doi:10.1016/j.neuron.2009.01.031.

[3] D. Weintraub, J. Koester, P. Mn, E. Al, Impulse control disorders in parkinson disease: A cross-sectional study of 3090 patients, Arch. Neurol. 67 (2010) 589–595.

[4] M. Brand, E. Kalbe, K. Labudda, E. Fujiwara, J. Kessler, H.J. Markowitsch, Decusion-making impairments in pations with Parkinson’s disease, Psychiatry Res. 133 (2005) 91–99. doi:10.1016/j.psychres.2004.10.003.

[5] F. Euteneuer, F. Schaefer, R. Stuermer, W. Boucsein, L. Timmermann, M.T. Barbe, G. Ebersbach, J. Otto, J. Kessler, E. Kalbe, Dissociation of decision-making under ambiguity and decision-making under risk in patients with Parkinson’s disease: A neuropsychological and psychophysiological study, Neuropsychologia. 47 (2009) 2882–2890. doi:10.1016/j.neuropsychologia.2009.06.014.

[6] A.C. Simioni, A. Dagher, L.K. Fellows, Dissecting the effects of disease and treatment on impulsivity in Parkinson’s disease, J. Int. Neuropsychol. Soc. 18 (2012) 942–951. doi:10.1017/S135561771200094X.

[7] A. Djamshidian, A. Jha, S.S. O’Sullivan, L. Silveira-Moriyama, C. Jacobson, P. Brown, A. Lees, B.B. Averbeck, Risk and learning in impulsive and nonimpulsive patients with Parkinson’s disease, Mov. Disord. 25 (2010) 2203–2210. doi:10.1002/mds.23247.

[8] S. Kobayashi, K. Asano, N. Matsuda, Y. Ugawa, Dopaminergic influences on risk preferences of Parkinson’s disease patients, Cogn. Affect. Behav. Neurosci. 19 (2018) 88–97. doi:10.3758/s13415-018-00646-3.

[9] M.H.M. Timmer, G. Sescousse, R.A.J. Esselink, P. Piray, R. Cools, Mechanisms underlying dopamine-induced risky choice in parkinson’s disease with and without depression (history), Comput. Psychiatry. 2 (2018) 11–27. doi:10.1162/cpsy_a_00011.

[10] V. Voon, J. Gao, C. Brezing, M. Symmonds, V. Ekanayake, H. Fernandez, R.J. Dolan, M. Hallett, Dopamine agonists and risk: Impulse control disorders in Parkinson’s; Disease, Brain. 134 (2011) 1438–1446. doi:10.1093/brain/awr080.

[11] D.O. Claassen, W.P.M. Van Den Wildenberg, K.R. Ridderinkhof, C.K. Jessup, M.B. Harrison, G.F. Wooten, S.A. Wylie, The Risky Business of Dopamine Agonists in Parkinson Disease and Impulse Control Disorders, Behav. Neurosci. 125 (2011) 492–500. http://dx.doi.org/10.1037/a0023795.supp (accessed February 15, 2018).

[12] M.T. Buelow, L.L. Frakey, J. Grace, J.H. Friedman, The Contribution of Apathy and Increased Learning Trials to Risky Decision-Making in Parkinson’s Disease, Arch. Clin. Neuropsychol. 29 (2014) 100–109. doi:10.1093/arclin/act065.

[13] T. Van Eimeren, B. Ballanger, G. Pellecchia, M. Janis, A Trigger for Pathological Gambling in Parkinson’s, Neuropsychopharmacology. 34 (2009) 2758–2766. doi:10.1038/sj.npp.npp2009124.Dopamine.

[14] M.E. Sharp, J. Viswanathan, M.J. McKeown, S. Appel-Cresswell, A.J. Stoessl, J.J.S. Barton, Decisions under risk in Parkinson’s disease: Preserved evaluation of probability and magnitude, Neuropsychologia. 51 (2013) 2679–2689. doi:10.1016/j.neuropsychologia.2013.08.008.

[15] J.M. Wenzel, N.A. Rauscher, J.F. Cheer, E.B. Oleson, A role for phasic dopamine release within the nucleus accumbens in encoding aversion: A review of the neurochemical literature, ACS Chem. Neurosci. 6 (2015) 16–26. doi:10.1021/cn500255p.

[16] M.E. Sharp, J. Viswanathan, L.J. Lanyon, J.J.S. Barton, Sensitivity and bias in decisionmaking under risk: Evaluating the perception of reward, its probability and value, PLoS One. 7 (2012) e33460. doi:10.1371/journal.pone.0033460.

[17] Z.S. Nasreddine, N.A. Phillips, V. Bedirian, S. Charbonneau, V. Whitehead, I. Collin, J.L. Cummings, H. Chertkow, The Montreal Cognitive Assessment, MoCA: A Brief Screening Tool For Mild Cognitive Impairment, J. Am. Geriatr. Soc. 53 (2005) 695–699. doi:10.1111/j.1532-5415.2005.53221.x.

[18] A.T. Beck, C.H. Ward, M. Mendelson, J. Mock, J. Erbaugh, An inventory for measuring depression., Arch. Gen. Psychiatry. 4 (1961) 561–71. http://www.ncbi.nlm.nih.gov/pubmed/13688369 (accessed February 9, 2018).

[19] H. Wynne, Introducing the Canadian problem gambling index, Edmonton, AB Wynne Resources. (2003).

[20] D. Bates, M. Mächler, B. Bolker, S. Walker, Fitting Linear Mixed-Effects Models using lme4, J. Stat. Softw. 67 (2015) 1–48. doi:10.18637/jss.v067.i01.

[21] V.F. Reyna, C.J. Brainerd, Numeracy, ratio bias, and denominator neglect in judgments of risk and probability, Learn. Individ. Differ. 18 (2008) 89–107. doi:10.1016/j.lindif.2007.03.011.

[22] M.J. Frank, L.C. Seeberger, R.C. O’Reilly, By carrot or by stick: Cognitive reinforcement learning in Parkinsonism, Science. 306 (2004) 1940–1943. doi:10.1126/science.1102941.

[23] R.B. Rutledge, N. Skandali, P. Dayan, R.J. Dolan, Dopaminergic Modulation of Decision Making and Subjective Well-Being, J. Neurosci. 35 (2015) 9811–9822. doi:10.1523/JNEUROSCI.0702-15.2015.

[24] F. Rigoli, R.B. Rutledge, B. Chew, O.T. Ousdal, P. Dayan, R.J. Dolan, Dopamine increases a value-independent gambling propensity, Neuropsychopharmacology. 41 (2016) 2658–2667. doi:10.1038/npp.2016.68.

[25] M.H.M. Timmer, G. Sescousse, R.A.J. Esselink, P. Piray, R. Cools, Mechanisms Underlying Dopamine-Induced Risky Choice in Parkinson ‘ s Disease With and Without Depression (History), Comput. Psychiatry. 2 (2018) 11–27. doi:10.1162/cpsy_a_00011.

[26] F. Niccolini, P. Su, M. Politis, Dopamine receptor mapping with PET imaging in Parkinson’s disease, J. Neurol. 261 (2014) 2251–2263. doi:10.1007/s00415-014-7302-2.

[27] A.G.E. Collins, M.J. Frank, Opponent actor learning (OpAL): Modeling interactive effects of striatal dopamine on reinforcement learning and choice incentive, Psychol. Rev. 121 (2014) 337–366. doi:10.1037/a0037015.

[28] C.J. Burke, A. Soutschek, S. Weber, A.R. Beharelle, E. Fehr, H. Haker, P.N. Tobler, Dopamine Receptor-Specific Contributions to the Computation of Value, (2017). doi:10.1038/npp.2017.302.

[29] D. Weintraub, E. Mamikonyan, K. Papay, J.A. Shea, S.X. Xie, A. Siderowf. Questionnaire for Impulsive-Compulsive Disorders in Parkinson’s Disease-Rating Scale. Mov Disord. 2012;27(2):242–7.

[30] C.R. Cloninger, T.R. Przybeck, D.M. Svrakic, R.D. Wetzel, R. Cloninger, R.D.P.E. Wetzel. The Temperament and Character Inventory (TCI)□: A Guide to Its Development and Use. First Edit. St. Louis, Missouri: Center for Psychobiology and Personality; 1994.

