## Supplement for "Dopamine replacement remediates risk aversion in Parkinson’s disease in a value-independent manner"

**Vancouver Ambiguity Task**

An additional Vancouver Ambiguity Task was administered to assess decision making under ambiguity as part of a larger study. The structure of the task resembled that of Vancouver Gambling Task, except that while one of the prospects had an explicitly stated probability and magnitude of gain, no information was given about the other prospect represented as a gift box (Figure 1B). There were 15 different combinations of magnitude and probability for the known prospects, each associated with an expected value. Unbeknownst to the participants, the same 15 combinations were used for the unknown prospect, but randomly and independent of the known prospect characteristics. There were 14 repetitions of each prospect for a total of 210 trials.

The Ambiguity Task was analyzed in the same way as the Vancouver Gambling Task (see main text), but with the models predicting the likelihood of choosing the known vs. the unknown prospect and the characteristics of only the known prospect considered as predictors.

**Results:**

There was no main effect of group (p=0.81) or interaction between group and medication state (p = 0.13). There was a main effect of “medication” state collapsing between patients and controls (b = 0.26, SE = 0.11, z= 2.24, p = 0.02), but this effect is difficult to interpret, as it was driven by a test order effect in the controls (b = 0.26, SE = 0.12, z= 2.15, p = 0.03), which was not due to a practice effect (p = 0.94). There were no significant effects of medication on its own or in interaction with expected value in the patients (p≥0.1), although there was a significant interaction of medication with the known prospect’ probability, such that probability had a greater influence on choice in the OFF than in the ON state (b = 0.26, SE = 0.12, z= 2.15, p = 0.03). However, overall the choice of PD patients was more driven by the expected value of the known gamble in both the medicated and the unmedicated state (group x EV interaction collapsing across medication states: b = 0.97, SE = 0.12, z=7.77, p < 0.0005). The choices made by the PD group were more sensitive both to the known prospect’s probability (group x probability interaction: b = 1.86.97, SE = 0.29, z=6.39, p < 0.0005) and to its magnitude (group x magnitude interaction: b = 0.11, SE = 0.03, z=3.44, p < 0.0006).

In summary, PD patients did not differ from controls in terms of their attitudes towards the unknown prospects. However, their choices were more strongly determined by the characteristics of the known prospect, a tendency that was more apparent in the OFF state.

**Feedback history effects:**

*Analyses*

We performed exploratory analyses of the effects of feedback history on choice in Parkinson’s patients and controls. These models examined the effects of trial feedback history (number of continuous winning, non-winning or losing outcomes prior to a given trial) on choice in interaction with group and medication. This was done using the lme4 package in R, using linear mixed effects models (glmer function) with a logistic link, as described in the main text of the manuscript.

*Results*

On the Vancouver Gambling Task, the length of a winning streak leading up to a trial predicted more risk-averse choice in both PD patients and controls (b = 0.43, SE = 0.03, z=16.48, p < 0.0005). There was also an interaction of winning streak length with group (b = 0.09, SE = 0.04, z=2.52, p < 0.01): longer winning streaks promoted risk aversion in controls more than in the patients. Conversely, similar to gambler’s fallacy, longer non-winning streaks predicted risker choices in both controls and patients (b = 0.58, SE = 0.03, z=16.63, p < 0.0005) without an interaction with group. These effects were independent of medication (ps ≥ 0.17).

Outcome history did not significantly influence choice in the Ambiguity task in either group (ps ≥ 0.14).

On the Vancouver Roulette Task, longer winning streaks increased the likelihood of accepting gambles (b = 20.48, SE = 7.35, z=2.84, p = 0.005) and of betting larger amounts (≥2 tokens: b = 1.28, SE = 0.05, z=28.18, p < 0.0005; 3 tokens: b = 0.92, SE = 0.04, z=22.40 p < 0.0005). While evident in both groups, this effect was less pronounced for the PD patients for accepting gambles (group x number of consecutive wins interaction: b = 18.34, SE = 7.35, z=2.5, p = 0.01), but more pronounced in the patients for betting larger amounts (≥2 tokens: b = 0.14, SE = 0.06, z=2.17, p = 0.03; 3 tokens: b = 0.28, SE = 0.06, z=4.71, p < 0.0005). Losing streaks had differential effects on betting in the PD patients versus controls. Longer losing streaks increased the likelihood of betting 2 or more tokens in the controls (b = 0.21, SE = 0.02, z=8.45, p <0.0005), but not in the patients (p=0.64). On the contrary, in the patients, longer losing streaks reduced the likelihood of placing the largest bet (b = 0.19, SE = 0.03, z=-5.97, p < 0.0005). Thus, while we observed an effect similar to gambler’s fallacy in the controls, there was no such effect in the patients; in fact it was reversed for betting the largest amount. Medication did not interact with the feedback history effects in the patients (ps ≥ 0.53).

Given our findings that choices of the Parkinson’s patients were more driven by the expected values of the prospects, we were interested in whether the weaker influence of expected value in controls may be partly attributed to their greater susceptibility to feedback history effects. We therefore examined whether feedback history interacted with EVR to predict choice, and whether this differed as a function of group. On the Vancouver Gambling Task, winning streaks moderated the effect of EVR on choice, such that longer winning streaks were associated with stronger effects of expected value (b = 0.32, SE = 0.06, z=5.78, p < 0.0005), however this effect did not interact with group (p=0.32), and the significant interaction of group with EVR remained in the presence of the trial history term (b = 0.47, SE = 0.12, z=3.79, p = 0.0001). Conversely, longer non-winning streaks were associated with weaker effects of expected value (b = 0.19, SE = 0.06, z=3.11, p = 0.002), but again, there were no interactions with group (ps ≥ 0.14). A similar effect was observed for the Ambiguity task, where longer non-winning streaks were likewise associated with weaker effects of expected value (b = 0.04, SE = 0.02, z=2.41, p = 0.02).

Because we did not observe group differences in the influence of expected value on gamble acceptance in the Roulette Task, the analysis focused on predicting the likelihood of betting 2 or more tokens. Across both groups, longer winning streaks increased the otherwise low likelihood of betting 2 or more tokens when those bets were associated with less favorable expected values (winning streak x EVR interaction: b = 0.2, SE = 0.05, z=4.52, p < 0.0005). A similar interaction was evident when considering 3-token bets (b = 0.19, SE = 0.03, z=7.17, p < 0.0005). For betting 2 or more tokens, this interaction differed as a function of group (winning streak x EVR x group interaction: b = 0.28, SE = 0.07, z=3.97, p < 0.0005) and was driven by the controls (winning streak x EVR interaction: b = 0.2, SE = 0.04, z=4.49, p < 0.0005). In the patients, longer winning streaks increased the likelihood of higher bets regardless of the expected value of placing those bets (main effect of winning streak: b = 0.83, SE = 0.05, z=16.01, p < 0.0005). Longer losing streaks promoted irrational choice, increasing the likelihood of betting 2 or more tokens when those bets were associated with less favorable expected values, while decreasing the likelihood of placing the larger bets when these bets were associated with higher expected values (≥ 2 tokens: b = 0.25, SE = 0.03, z=9.04, p < 0.0005; 3 tokens: b = 0.11, SE = 0.02, z=6.33, p < 0.0005), but this did not differ as a function of group (ps≥0.47). Even in the presence of a significant interaction with group in the case of winning streaks, these feedback history effects did not fully account for the increased influence of expected value on choice in the patients: the significant interaction between group and expected value remained with the winning streak term present in the model (b = 0.13, SE = 0.06, z=2.2, p < 0.03). Thus, although feedback history had significant effects on choice in both the patients and the controls, with some suggestion of weaker influence on the patients, this did not account for the greater value-dependence of the patients’ choices.
